# Cross-subject brain entropy mapping

**DOI:** 10.1101/2024.04.05.588307

**Authors:** G. Del Mauro, Z. Wang

## Abstract

We present a method to map the regional similarity between resting state fMRI activities of different individuals. The similarity was measured using cross-entropy. Group level patterns were displayed based on the Human Connectome Project Youth data. While we only showed the cross-subject brain entropy (BEN) mapping results in this manuscript, the same concept can be directly extended to map the cross-sessional BEN and the cross-regional cross-subject or subject-session BEN.

## Introduction

Analyzing brain structural and functional properties to identify general patterns and draw population-level inferences has been a major focus of human neuroimaging research. Human brain, however, displays significant inter-individual differences in both its structural (1–4) and functional (5–7) facets, which underlie the complex and astonishing variability of human behavior. In recent years, scientific community in cognitive neuroscience has prompted a shifting from population-to individual-level inferences, aiming at studying individual brain fingerprints to aid predicting unique individual traits. In particular, recent findings suggest that brain functional-connectivity (FC) patterns can be adopted to predict cognition, personality traits, emotions as well as individuals’ identity (5, 8–10).

Brain entropy (BEN) has been increasingly used to investigate the temporal coherence and variability of brain activity. In fact, BEN measures the repeatability over time of the patterns of blood-oxygen-level-dependent (BOLD) signal, and as such represents an indirect index of the brain activity temporal coherence: higher BEN indicates lower temporal coherence (i.e., higher randomness), while lower BEN indicates higher temporal coherence (i.e., lower randomness) of brain signal (11). BEN, and in particular resting-state fMRI (rs-fMRI) based BEN, has been shown to correlate with both biological (e.g., age and sex) and cognitive variables (e.g., fluid intelligence) (12, 13). While BEN as described above allows to draw population-level inference, it does not enable however to measure the degree of similarity between one subject’s brain activity fingerprint to another’s. In this pilot study based on the Human Connectome Project – Young Adult (HCP-Y) dataset we propose a variation of BEN, namely cross-subject (cs-BEN), to aid investigating between-subjects similarity of brain activity. The significance of this new extension is that we can for the first time directly compare brain activity across different scan time or across different individuals. Related research is the movie-watching based inter-subject correlation analysis which however depends on the use of specific task paradigm such as movie-watching, and the correlation results may be subject to the large inter-subject variability of activity onset and magnitude.

## Materials and Methods

### Data

Fully pre-processed rs-fMRI data were downloaded from 1096 healthy young participants (mean age = 28.83 ± 3.69 years; mean education = 14.86 ± 1.81 years; males/females = 550/656) of the HCP-Y dataset. The HCP-Y protocol includes the acquisition of four rs-fMRI runs collected over two days. Each day involved the acquisition of a rs-fMRI session (Rest1 and Rest2) composed of two 15-minutes open-eyes runs. Each run was acquired using the same multi-band sequence and included 1200 timepoints. To compensate for image distortions induced by the long scan time, readout direction was from left to right for the first run of each session (Rest1 LR and Rest2 LR), and from right to left for the second run of each session (Rest1 RL and Rest2 RL). Other acquisition parameters included: repetition time (TR) = 720ms, echo time (TE) = 33.1ms, resolution = 2x2x2mm^3^. (see (14) for detailed information regarding pre-processing pipeline).

### Cross-entropy (CrEn) cross-subject BEN (cs-BEN)

cs-BEN was estimated using cross-entropy (CrEn), defined as the distance of the probability distributions of two random variables. For cross-subject entropy mapping, the two variables are two time series, each from a different individual subject. To avoid the issue of unstable probability density estimation for short time series, in this study we used the cross-sample entropy (SampEn) as the approximate formula for calculating CrEn. Similar to SampEn, cross-SampEn is based on temporal data segment matching, but the matching crosses different time series rather than within the same time series. The CrEn estimation pipeline is described in **Fig. 1**. While this process is nearly identical to SampEn calculation, except for the cross-time series segment extraction and matching (there is no within time series segment matching), the total number of segments is doubled and therefore the number of segments compared is more than doubled as compared to SampEn. To mitigate the large computational burden, we implemented the algorithm using C++ and parallel computing. The difference between CrEn and the widely used correlation coefficient (CC)-based FC is showed in **Fig. 2**, which clearly illustrates the incapability of the CC-based FC to capture the cross-time inter-areal or inter-nodal interactions or information exchange.

**Fig. 1.**
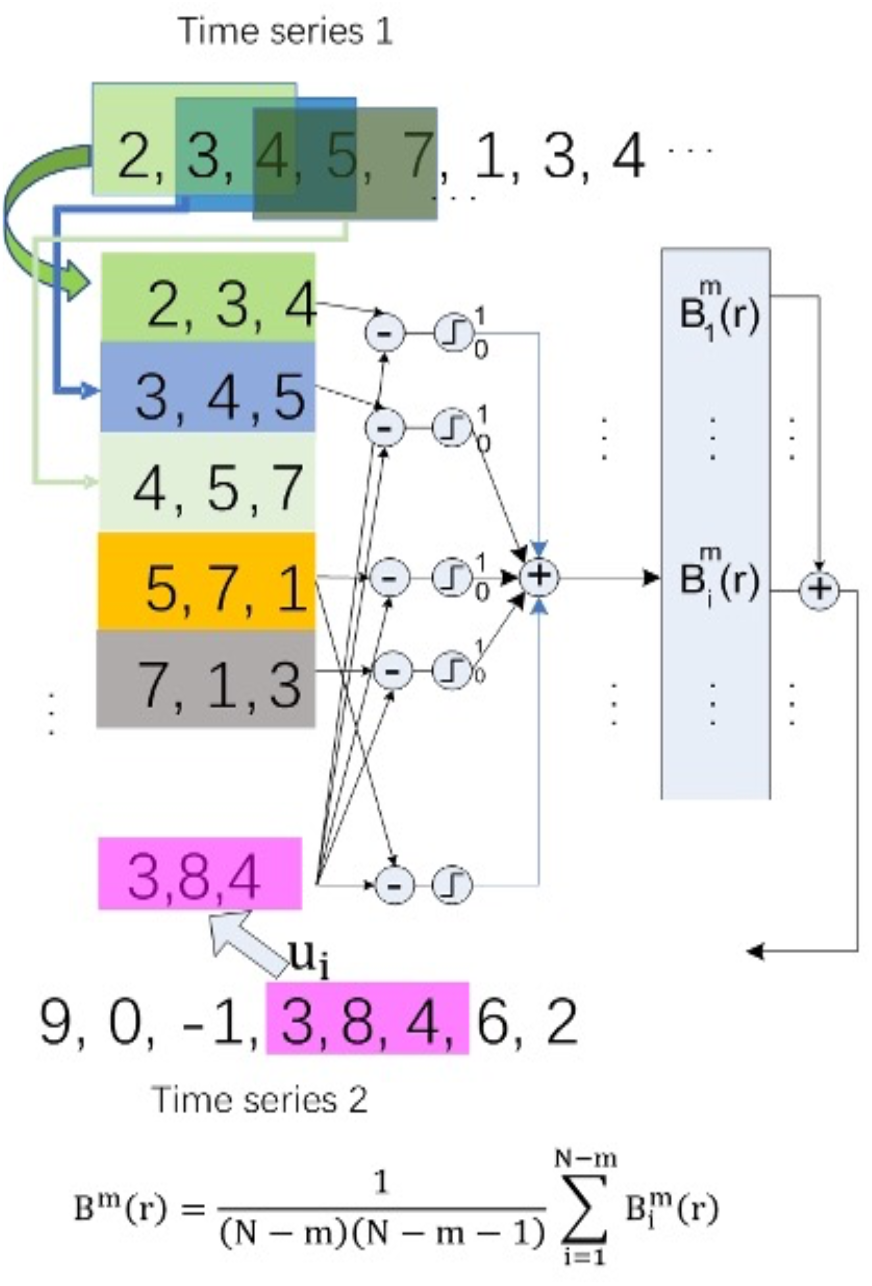
The data matching process in Cross-Entropy (CrEn) calculation. For an arbitrary segment u_i_ extracted from Time Series 2 (purple box), it will be compared to all temporal segments of Time Series 1. The total number of matches will be recorded by 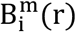. The sum of total matches of all temporal segments of Time Series 2 (or exchangeable of Time Series 1) will be counted as 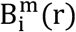. A similar process can be performed to get 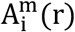 when the observation window is increased by 1. The CrEn can be then calculated as –ln 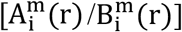.

**Fig. 2.**
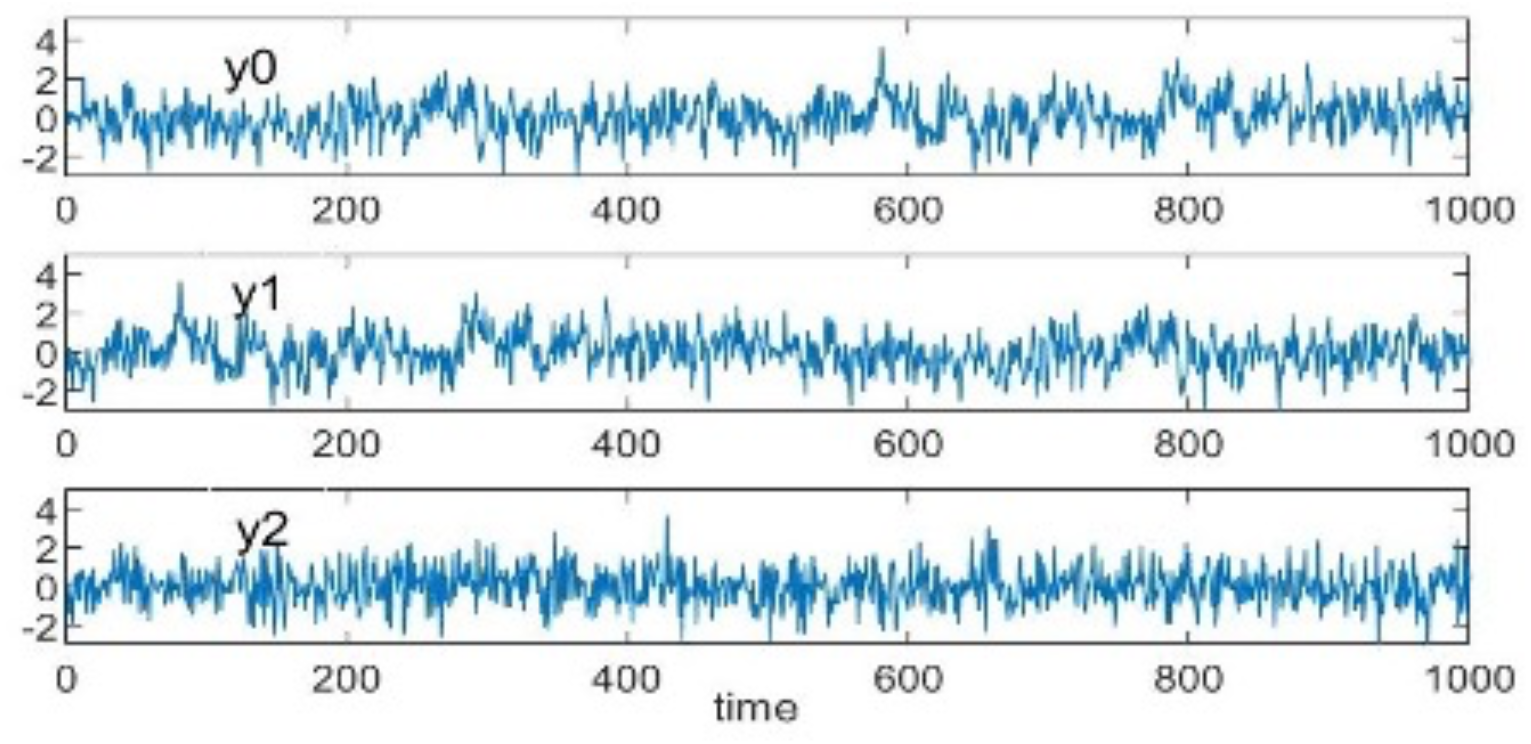
A power noise (y0), its half shifted version (y1), and the randomly time shuffled version (y2). To illustrate the difference between cross-entropy (CrEn) and functional connectivity (FC), we generated three long time series: y0, y1, and y2 (the time series are showed in **Fig. 1** and differ only by time positions). Using the cross-SampEn as the approximation to CrEn, we get CrEn(y0, y1) = 1.70, which is the same as CrEn(y0, y0) =1.70 (the same as the entropy of y0). Even after a random position shuffling (y2) CrEn is still very close to the ground truth (CrEn(y0, y2)=1.75) (the moderate offset is likely due to the imperfection of the cross-SampleEn approximation for CrEn). By contrast, correlation coefficient (CC) of (y0, y1) becomes -0.01 and CC(y0, y2) becomes 0.01, even though the only change is the position of the data points.

In this study, CrEn was used to calculate the cs-BEN maps for the HCP-Y dataset. The calculation was performed for each of the four rs-fMRI scans separately. For each scan, each subject was paired with any other subject to calculate a cs-BEN map. The value of cs-BEN at each voxel was calculated using the time series from the corresponding two subjects at that voxel. For each rs-fMRI scan and each participant, we obtained: 1074 cs-BEN maps for Rest1 LR, 1088 for Rest1 RL, 994 for Rest2 LR, and 1008 for Rest2 RL (note that not all participants completed four rs-fMRI runs). Then, for each subject we calculated a mean and standard deviation (std) map using the cs-BEN maps between that subject and all others.

This process was followed for each rs-fMRI scan separately. In the end, for each scan, a mean and a std cs-BEN map was obtained from each participant. Finally, mean and std cs-BEN maps were converted to Z-scores and smoothed with a 4x4x4 mm full width at half maximum (FWHM) filter before statistical analyses.

### Statistical analysis

To investigate which brain regions had significantly high/low cs-BEN as well as high/low cs-BEN variability, a one-sample t-test was performed for each run separately on the mean and std cs-BEN maps of each participant. Results were corrected for multiple comparisons at the voxel level using the family-wise error (FWE) methods and considered significant if p-FWE < 0.05.

## Results

One-sample t-tests on the mean cs-BEN maps resulted in significantly lower cs-BEN in the frontal, parietal, and occipital lobes, while regions with higher cs-BEN included the temporal lobes, subcortical areas, cerebellum, and white matter (**Fig. 3**). Statistical analyses performed on the std cs-BEN maps resulted in a reversed pattern, with frontal, parietal, and occipital lobes showing significantly higher standard deviation, and temporal lobes, subcortical regions, cerebellum, and white matter displaying lower standard deviation (**Fig. 4**).

**Fig. 3.**
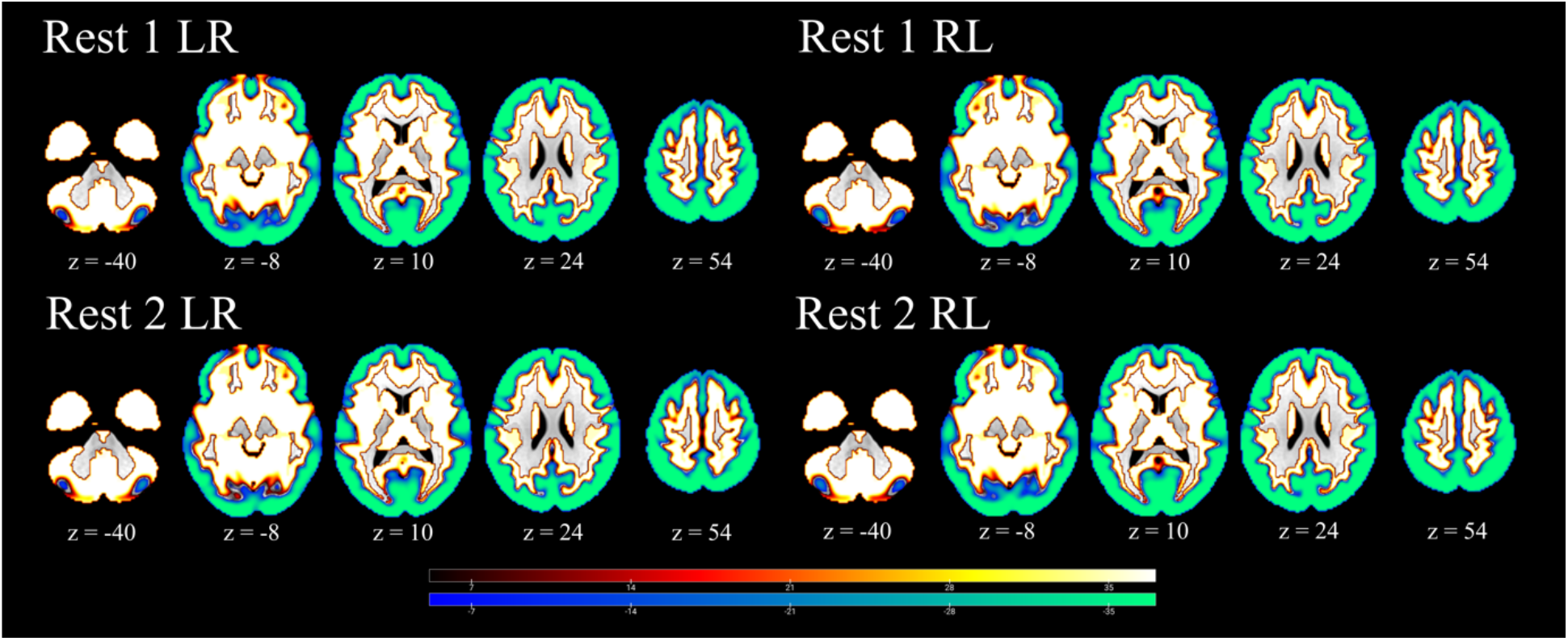
One-sample t-tests performed on the mean cross-subject brain entropy (cs-BEN) maps. Analyses were performed on four resting-state fMRI (rs-fMRI) separately. Results were corrected for multiple comparisons at the voxel level using the family-wise error (FWE) method. Results were considered significant if p-FWE < 0.05.

**Fig. 4.**
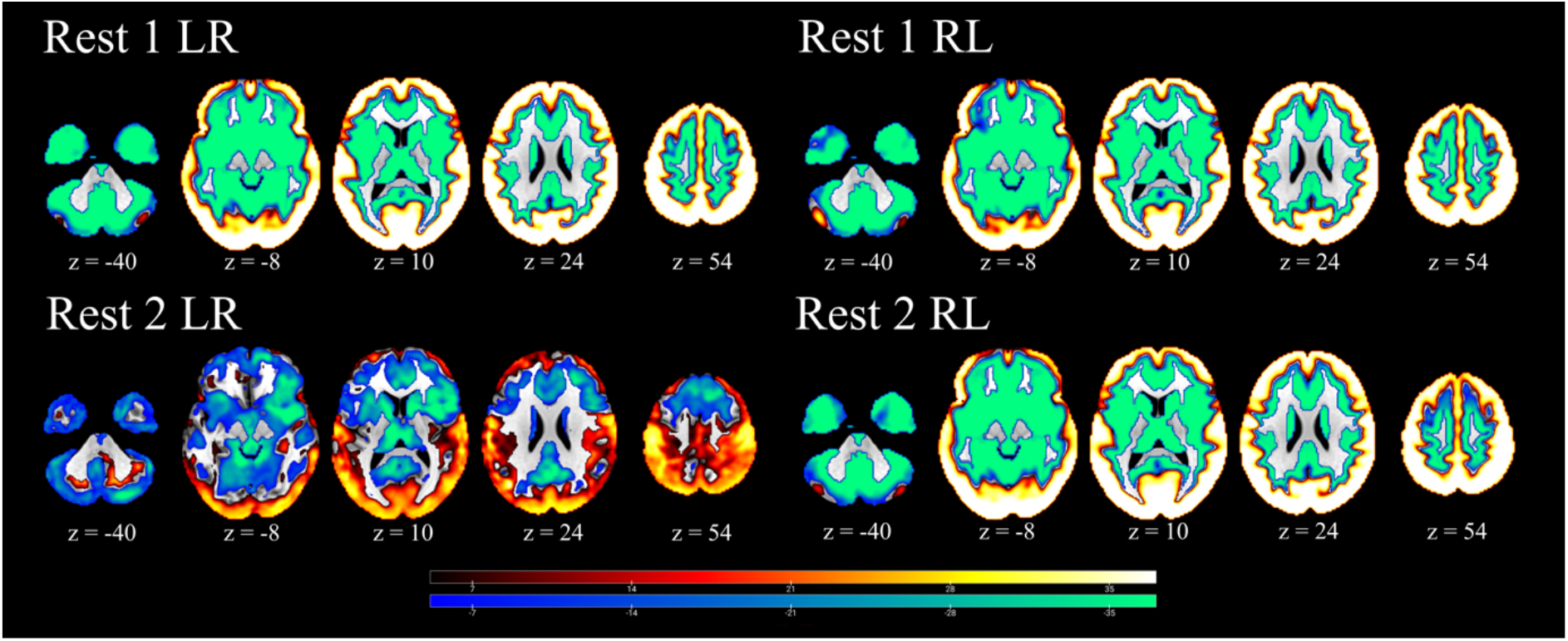
One-sample t-tests performed on the standard deviation (std) cross-subject brain entropy (cs-BEN) maps. Analyses were performed on four resting-state fMRI (rs-fMRI) separately. Results were corrected for multiple comparisons at the voxel level using the family-wise error (FWE) method. Results were considered significant if p-FWE < 0.05.

## Discussion

In this work, we for the first time measured similarity of rs-fMRI data between different individuals’ brains by computing the cross entropy. In this context, higher cross-subject brain entropy (cs-BEN) reflects lower degree of similarity, and lower cs-BEN a higher degree of similarity between the BOLD time series of two subjects. For each participant, distinct cs-BEN maps were calculated between that participant and all others. Finally, all cs-BEN maps of the same participant were used to generate a mean and a std cs-BEN map. This process was followed for four distinct rs-fMRI runs. The results of the one-sample tests performed on the mean and std cs-BEN maps were highly consistent across runs. In particular, we showed that the activity of frontal, parietal, and occipital lobes consistently displayed a lower mean and a larger standard deviation of the cs-BEN. This result suggests that, despite showing a high similarity of the BOLD time series (i.e., lower mean), there is a large variability in how the activity of these regions is similar in pairs of individuals (i.e., higher standard deviation). On the other hand, other regions including temporal lobes, subcortical areas, and cerebellum showed the inverse pattern, corresponding to low similarity of the BOLD signal (i.e., higher mean of cs-BEN) and a narrowed range of values in pairs of individuals (i.e., lower standard deviation of cs-BEN). Future studies may investigate the relationship between cs-BEN and biological (e.g., age and sex) as well as cognitive variables.

While we only showed cross-subject BEN mapping results, the same concept can be directly extended to a cross-sessional BEN mapping or a cross-subject (or cross-session) and cross regional BEN mapping.

## Acknowledgement

This work was supported by the National Institute on Aging [R21AG082345, R01AG060054, R01AG070227, R01AG081693] and the University of Maryland Baltimore, Institute for Clinical & Translational Research (ICTR) [1UL1TR003098].

